# MegaGO: a fast yet powerful approach to assess functional similarity across meta-omics data sets

**DOI:** 10.1101/2020.11.16.384834

**Authors:** Pieter Verschaffelt, Tim Van Den Bossche, Wassim Gabriel, Michał Burdukiewicz, Alessio Soggiu, Lennart Martens, Bernhard Y. Renard, Henning Schiebenhoefer, Bart Mesuere

## Abstract

The study of microbiomes has gained in importance over the past few years, and has led to the fields of metagenomics, metatranscriptomics and metaproteomics. While initially focused on the study of biodiversity within these communities the emphasis has increasingly shifted to the study of (changes in) the complete set of functions available in these communities. A key tool to study this functional complement of a microbiome is Gene Ontology (GO) term analysis. However, comparing large sets of GO terms is not an easy task due to the deeply branched nature of GO, which limits the utility of exact term matching. To solve this problem, we here present MegaGO, a user-friendly tool that relies on semantic similarity between GO terms to compute functional similarity between two data sets. MegaGO is highly performant: each set can contain thousands of GO terms, and results are calculated in a matter of seconds. MegaGO is available as a web application at https://megago.ugent.be and installable via pip as a standalone command line tool and reusable software library. All code is open source under the MIT license, and is available at https://github.com/MEGA-GO/.

## Introduction

Microorganisms often live together in a microbial community or microbiome, where they create complex functional networks. These microbiomes are therefore commonly studied to reveal both their taxonomic composition as well as their functional repertoire. This is typically achieved by analyzing their gene content using shotgun metagenomics. While this approach allows a quite detailed investigation of the genomes that are present in such multi-organism samples, it only reveals their functional potential rather than their currently active functions^1^. To uncover these active functions within a given sample, the characterization of the protein content is often essential^2^.

The growing focus on functional information as a complement to taxonomic information^3^ is derived from the observation that two taxonomically similar microbial communities could have vastly different functional capacities, while taxonomically quite distinct communities could have remarkably similar functions. While the investigation of the active functions is thus increasingly seen as vital to a complete understanding of a microbiome, the identification and comparison of these detected functions remains one of the biggest challenges in the field^4^.

Several omics tools exist to describe functions in microbial samples, although these tools link functionality to different biological entities such as genes, transcripts, proteins and peptides^5-14^. However, most tools are capable of directly or indirectly reporting functional annotations as a set of Gene Ontology^15^ (GO) terms, regardless of the biological entity it is assigned to. In October 2020, there exist 44 264 of these terms in the complete GO tree. GO terms are organized in three independent domains: molecular function, biological process, and cellular component^16^. In each domain, terms are linked into a directed acyclic graph, an excerpt of which is shown in **Figure 1**. In the GO graph, a parent term can have one or more children (e.g., the root node “biological process” is the parent of the children GO:0009987 and GO:0008152), and children can have multiple parents (e.g., the most specific term “translation” has as parents GO:0043043, GO:0034645 and GO:0044267).

**Figure 1.**
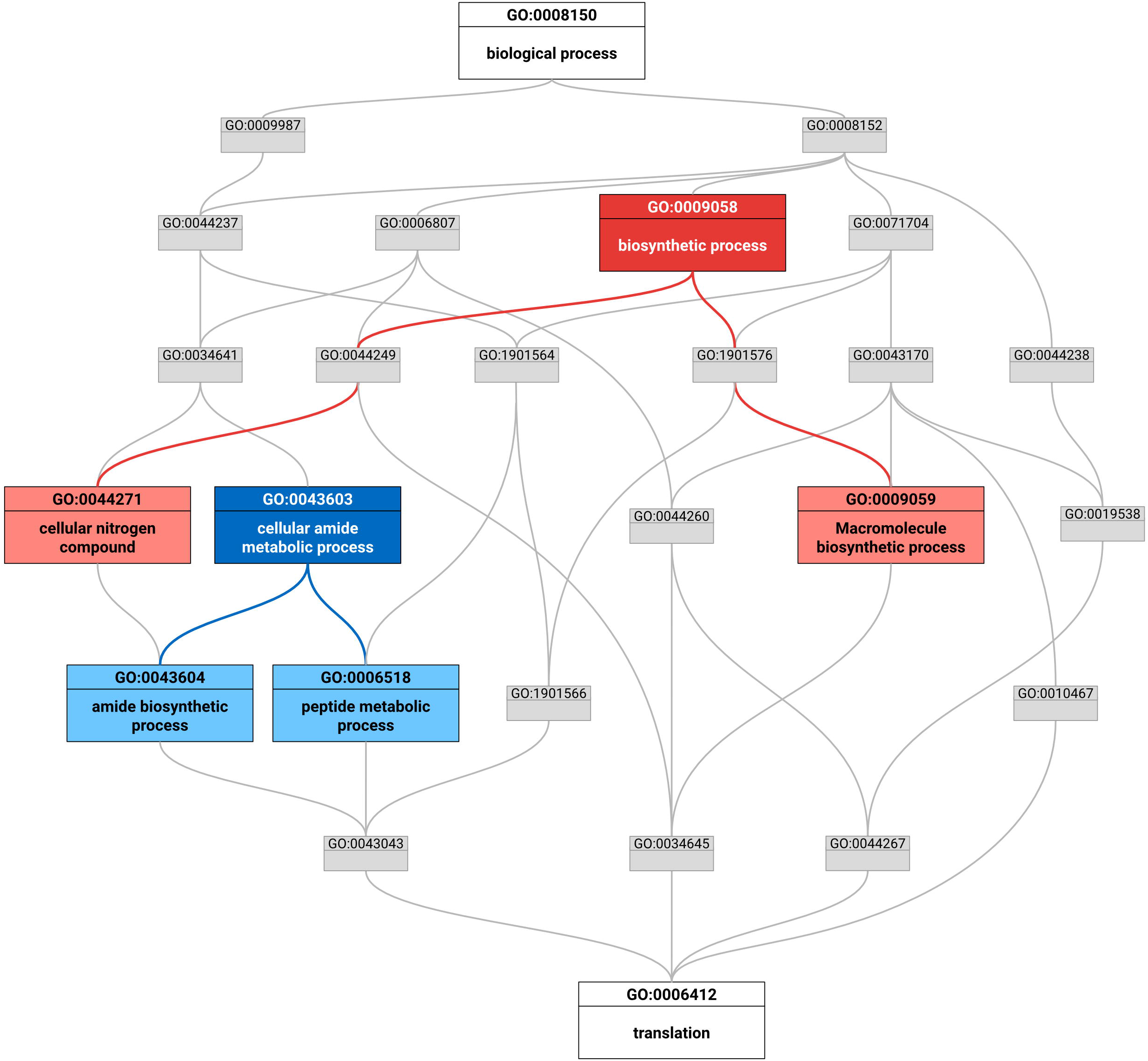
Excerpt of the biological process domain of the Gene Ontology showing all parent terms up to the root for “translation” (GO:0006412). The root GO term “biological process” (GO:0008150) has multiple children. The most specific term “translation”, in contrast, has multiple parents. When comparing the two terms GO:0044267 and GO:0034645 (portrayed in light red), we find two different lowest common ancestors: GO:0044249 and GO:1901576 (dark red). Only one of these, however, can be the most informative common ancestor (MICA), i.e. the common ancestor with the highest information content for the terms in light red. Since 1.52 is larger than 1.48, the GO:0044249 is the MICA. The terms GO:0043604 and GO:0006518 (in light blue) are more similar than the two terms we described earlier and only have one lowest common ancestor, which is also automatically the MICA for these terms: GO:0043603 (in dark blue). IC: information content, star: most informative common ancestor.

While this highly branched graph structure of GO allows flexible annotation at various levels of detail, it also creates problems when the results from one data set are compared to another data set. Indeed, even though two terms may be closely linked in the GO tree and are therefore highly similar (e.g., as parent and child terms, or as sibling terms), typically employed exact term matching will treat these terms as wholly unrelated as the actual GO terms (and their accession numbers) are not identical. This problem is illustrated in a study by Sajulga et al.^17^, where a multi-sample data set was analyzed using several metaproteomics tools. The resulting GO terms were then compared using exact matching. The overlap between the result sets was quantified using the Jaccard Index and was found to be quite low. As explained above, this low similarity is likely the result of the limitations of the exact term matching approach.

There is thus a clear need for a more sophisticated GO term comparison that takes into account the existing relationships in the full GO tree. However, most existing tools that provide such comparison are based on enrichment analyses^18-20^. In such analyses, a list of genes is mapped to GO terms, which are then analyzed for enriched biological phenomena. As a result, to the best of our knowledge no tools allow the direct comparison of large functional data sets against each other, nor are these able to provide metrics to determine how functionally similar two data sets are.

We therefore present MegaGO, a tool for comparing the functional similarity between two large lists of GO terms. MegaGO calculates a similarity score between two sets of GO terms for each of the three GO domains, and can do so in seconds, even on platforms with limited computational capabilities.

## Implementation

In order to measure the similarity of two sets of GO terms, we first need to measure the similarity of two individual terms. We compare two terms using the Lin semantic similarity metric^21^, which can take on a value between 0 and 1 (**Supplementary Formula 1a**). The Lin semantic similarity is based on the ratio of the information content of the most informative common ancestor (MICA) to the average of the terms’ individual information content.

The information content (**Supplementary Formula 1b**) is computed by estimating the terms’ probability of occuring (**Supplementary Formula 1c**), including that of all of its children. Term frequencies are estimated based on the manually curated SwissProt database. As a result, a high level GO term such as “biological process” (through its many direct or indirect child terms) will be present in all data sets, and thus carries little information. A more specific term such as “translation” (or any of its potential child terms) will occur less frequently, and thus be more informative (**Figure 1**). To finally calculate the similarity of two terms, we compare their information content with that of their shared ancestor that has the highest information content, the MICA. If the information content of the MICA is similar to the term’s individual information content, the terms are deemed to be similar. The dissimilar terms “peptide biosynthetic process” and “cellular macromolecule biosynthetic” are situated further from their MICA “cellular biosynthetic process” than the similar terms “amide biosynthetic process” and “peptide metabolic process” with their respective MICA “cellular amide metabolic process” (**Figure 1**).

MegaGO, however, can not only compare two terms but also two lists of GO terms. To achieve this, pairwise term similarities are aggregated using the Best Matching Average (BMA, **Supplementary Formula 2**)^22^. For each GO term in the first input data set, BMA finds the GO term with the highest Lin semantic similarity in the second data set and averages the values of these best matches. Moreover, MegaGO calculates the similarity for each of the three domains of the gene ontology (molecular function, biological process, and cellular component), as GO terms from distinct domains do not share parent terms. The general overview of MegaGO is shown in **Figure 2**.

**Figure 2.**
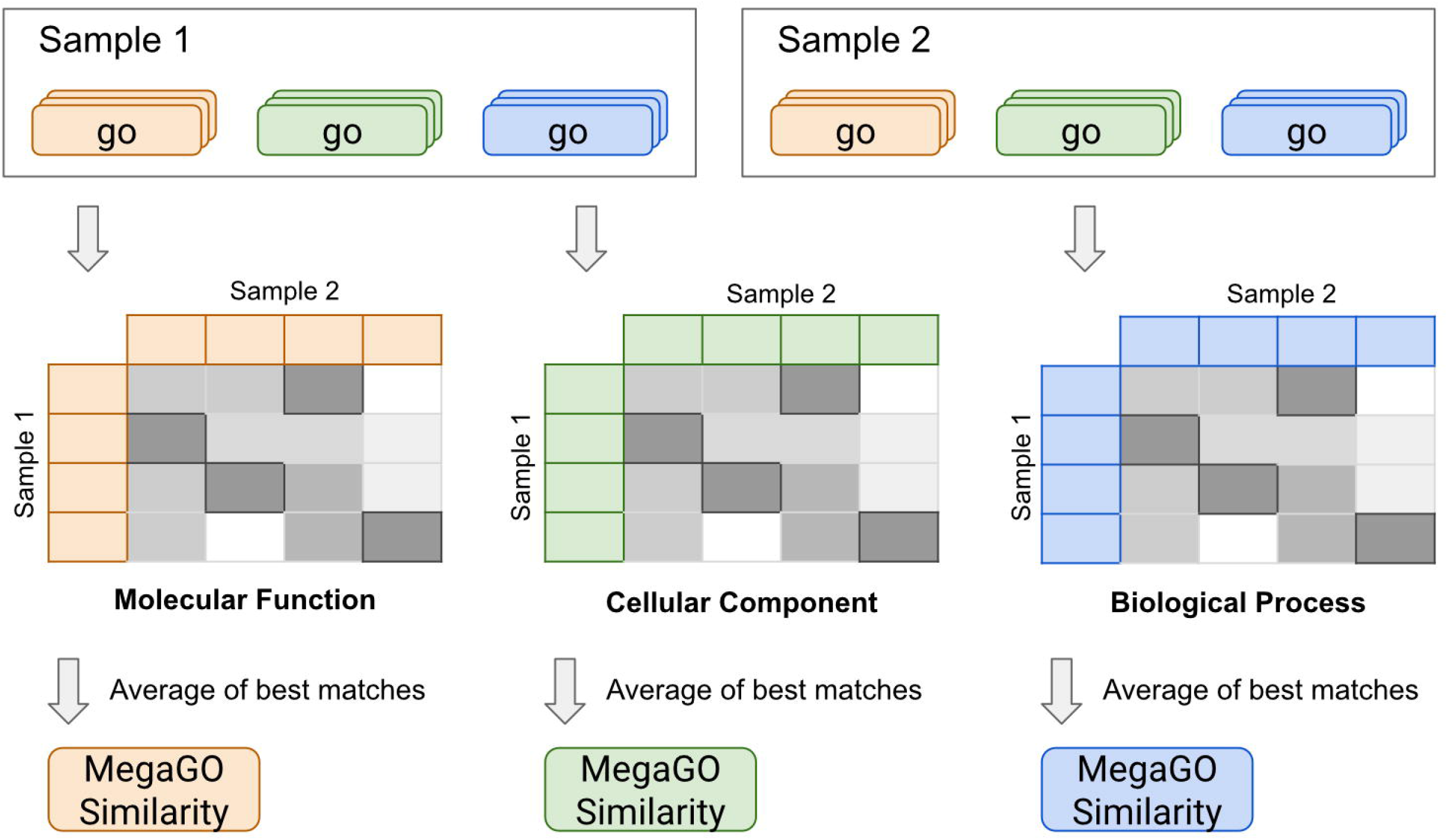
Overview of MegaGO workflow. Gene Ontology (GO) terms of each sample set are separated into three GO domains: molecular function, cellular component, and biological process. Each term of each sample set is compared to every term in the other set that is from the same domain. The match with highest similarity for each term is then selected and the average across all these best matches is calculated.

MegaGO is implemented in Python and installable as a Python package from PyPi and can easily be invoked from the command line. The GOATOOLS^23^ library is used to read and process the Gene Ontology and to compute the most informative common ancestor of two GO terms, which are both required to compute the information content value (**Supplementary Formula 1, p(go)**). GO-term counts are recomputed with every update of SwissProt and a new release is automatically published bi-monthly to PyPi which includes the new data set. Automated testing via GitHub Actions is in place to ensure correctness and reproducibility of the code. In addition, we also developed a user-friendly and easily accessible web application that is available on https://megago.ugent.be. The backend of the web application is developed with the Flask web framework for Python and the frontend uses Vue. Our web application has been tested on Chromium-based browsers (Chrome, Edge, Opera), as well as Mozilla Firefox and Safari. The MegaGO application is also available as a Docker-container on Docker Hub and can be started with a single click and without additional configuration requirements. Our Docker container is automatically updated at every change to the underlying MegaGO code. All code is freely available under the permissive open source MIT license on https://github.com/MEGA-GO/. Documentation for our Python script can be found on our website: https://megago.ugent.be/help/cli. A guide on how to use the web application is also available: https://megago.ugent.be/help/web.

MegaGO is cross-platform, and runs on Windows, macOS or Linux systems. Systems requirements are at least 4GiB of memory and support for either Python 3.6 (or above), or the Docker runtime.

## Validation

To validate MegaGO, we reprocessed the functional data from Easterly *et al*^24^. This data set consists of 12 paired oral microbiome samples that were cultivated in bioreactors. Each sample was treated with and without sucrose pulsing, hereafter named *ws* and *ns* samples, respectively. Samples were annotated with 1718 GO-terms on average. We calculated the pairwise similarity for each of the 300 sample combinations, which took less than a minute for a single sample pair on the web version of MegaGO. This resulted in a MegaGO similarity score for each of the three GO domains for each sample combination. These similarities were then hierarchically clustered and visualized in a heatmap. All data and intermediate steps of our data analysis are available at https://github.com/MEGA-GO/manuscript-data-analysis/.

In the heatmap (**Figure 3**) we can observe that the two sample groups cluster together, except for 730ns and 733ns that are clustered in the *ws* sample group. These two samples were also identified as outliers in Easterly *et al*^24^, and 733ns was originally also identified as both a taxonomic and functional outlier in Rudney *et al*^25^ Similar results can be observed for the GO domain ‘molecular function’ (**Supplementary Figure 1**). MegaGO similarity-based clustering of cellular component GO terms (**Supplementary Figure 2**) has two additional samples clustered outside of their treatment group: 852ws in the *ns* cluster and 861ns in the *ws* group. Again, these patterns can also be found in previous analyses: 852ws is placed in direct proximity of the ns samples in the PCA of HOMINGS analysis by Rudney *et al.*, 861ns is closest to 730 and 733ns in PCA of Rudney *et al.* ‘s taxonomic analysis. Interestingly, subjects 730 and 852 were the only ones without active carious lesions, which could cause their divergence in the similarity analyses.

**Figure 3.**
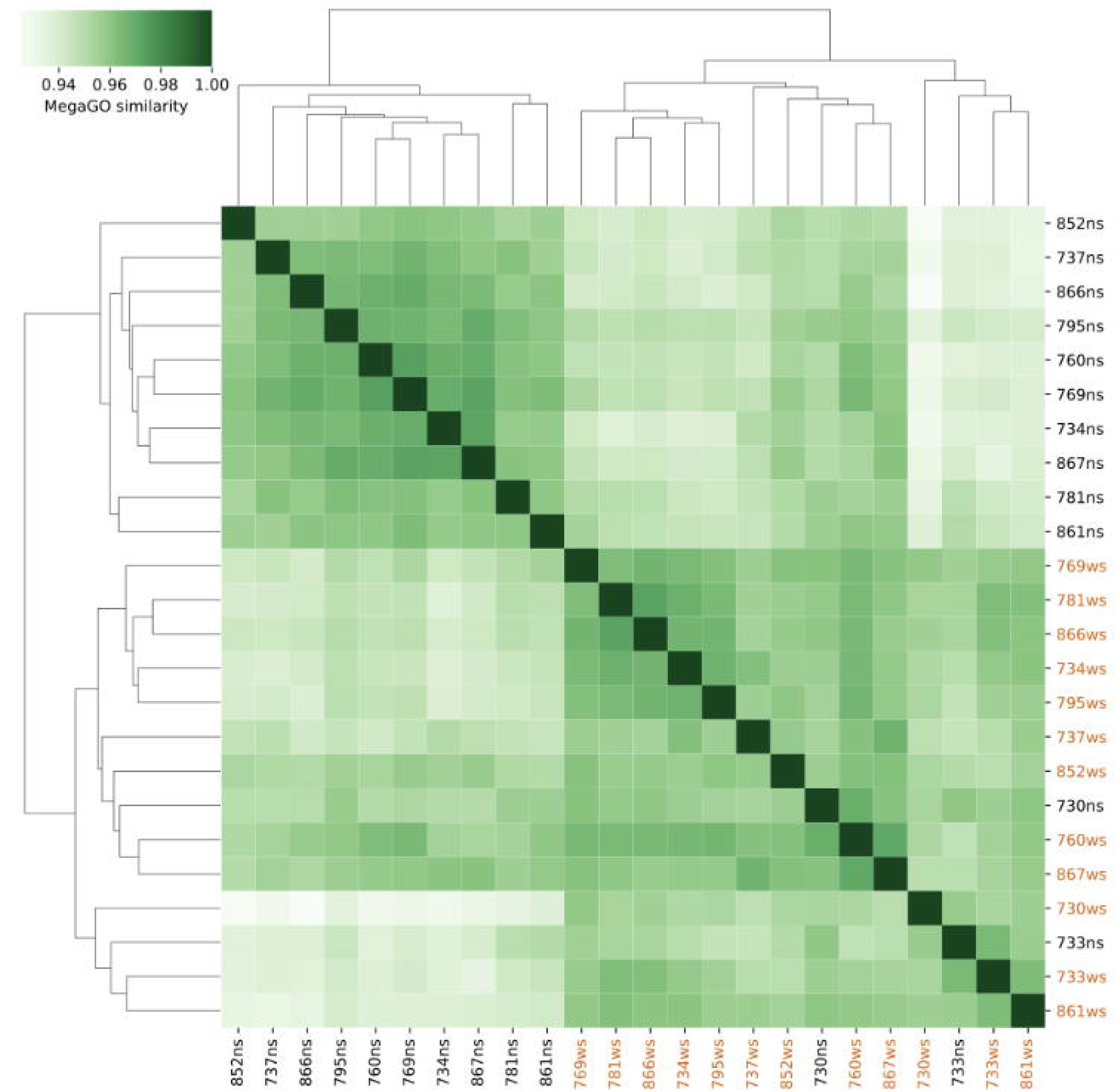
Hierarchically clustered heatmap comparing MegaGO similarities for the GO domain ‘biological process’ for each of the samples from Easterly *et al.*^24^. Samples that are treated with sucrose pulsing are labeled as “ws” and displayed in orange

Results produced by MegaGO are thus in close agreement with prior analyses of the same data, showing that MegaGO offers a valid and very fast approach for comparing the functional composition of samples.

## Conclusion

MegaGO enables the comparison of large sets of GO-terms, allowing users to efficiently evaluate multi-omics data sets containing thousands of terms. MegaGO calculates a similarity for each of the three GO-domains separately (biological process, molecular function, and cellular component). In the current version of MegaGO, quantitative data is not taken into account, thus giving each GO term identical importance in the data set.

MegaGO is compatible with any upstream tool that can provide GO term lists for a data set. Moreover, MegaGO allows the comparison of functional annotations derived from DNA, RNA, or protein based methods as well as combinations thereof.

## Supporting information

Supplemental formulas and figures

## Acknowledgements

We would like to acknowledge the European Bioinformatics Community for Mass Spectrometry (EuBIC-MS). This project found its origin at the EuBIC Developers’ 2020 meeting in Nyborg, Denmark. We would like to thank Thilo Muth and Stephan Fuchs for their support. PV, TVDB, LM and BM would like to acknowledge Research Foundation - Flanders (FWO) [grants 1164420N, 1S90918N, G042518N and 12I5220N]. LM also acknowledges support from the European Union’s Horizon 2020 Programme under Grant Agreement 823839 [H2020-INFRAIA-2018-1]. HS and BYR acknowledge support by Deutsche Forschungsgemeinschaft (DFG; grant number RE3474/5-1 and RE3474/2-2) and the BMBF-funded de.NBI Cloud within the German Network for Bioinformatics Infrastructure (de.NBI; 031A537B, 031A533A, 031A538A, 031A533B, 031A535A, 031A537C, 031A534A and 031A532B).

