## Supplemental formulas and figures for "MegaGO: a fast yet powerful approach to assess functional similarity across meta-omics data sets"

### Supplementary Formula 1

a) $sim_{Lin}(go_1, go_2) = \frac{IC(MICA)}{\frac{IC(go_1)+IC(go_2)}{2}}$

b) $IC(go_i) = -\, log(p(go_i))$

c) $p(go_i) = \frac{\sum_{go' \in \{go_i\} \cup c} n_{go'}}{N}$

**Formula 1**:

a) Lin semantic similarity (simLin) metric with *IC* as the information content, defined by b), and *MICA* as the Most Informative Common Ancestor of GO term 1 and GO term 2.

b) The information content requires the frequency *p(go_i)* of the i-th GO term as defined by c) and provides a notion for how likely an event is to occur randomly.

c) Defines the probability of a GO term $go_i$ , where $c$ is the set of all (direct and indirect) children of $go_i$ , $N$ is the total number of terms in the GO corpus and $n_{go'}$ is the number of occurrences of a term $go'$ in a reference data set.

### Supplementary Formula 2

$$SIM_{BMA}(g_1, g_2) = \frac{1}{m+n} * \left( \sum_{1=i}^{m} \max_{1 \le j \le n} (sim(go_{1i}, go_{2j})) + \sum_{1=j}^{n} \max_{1 \le i \le m} (sim(go_{1i}, go_{2j})) \right)$$

**Formula 2**: the best match average (BMA) to calculate the similarity of two sets of terms, with *m,n* as the number of terms in set $g_i$ and $g_j$, respectively, and $sim(go_{1i}, go_{2j})$ as the similarity between two GO terms.

### Supplementary Figure 1

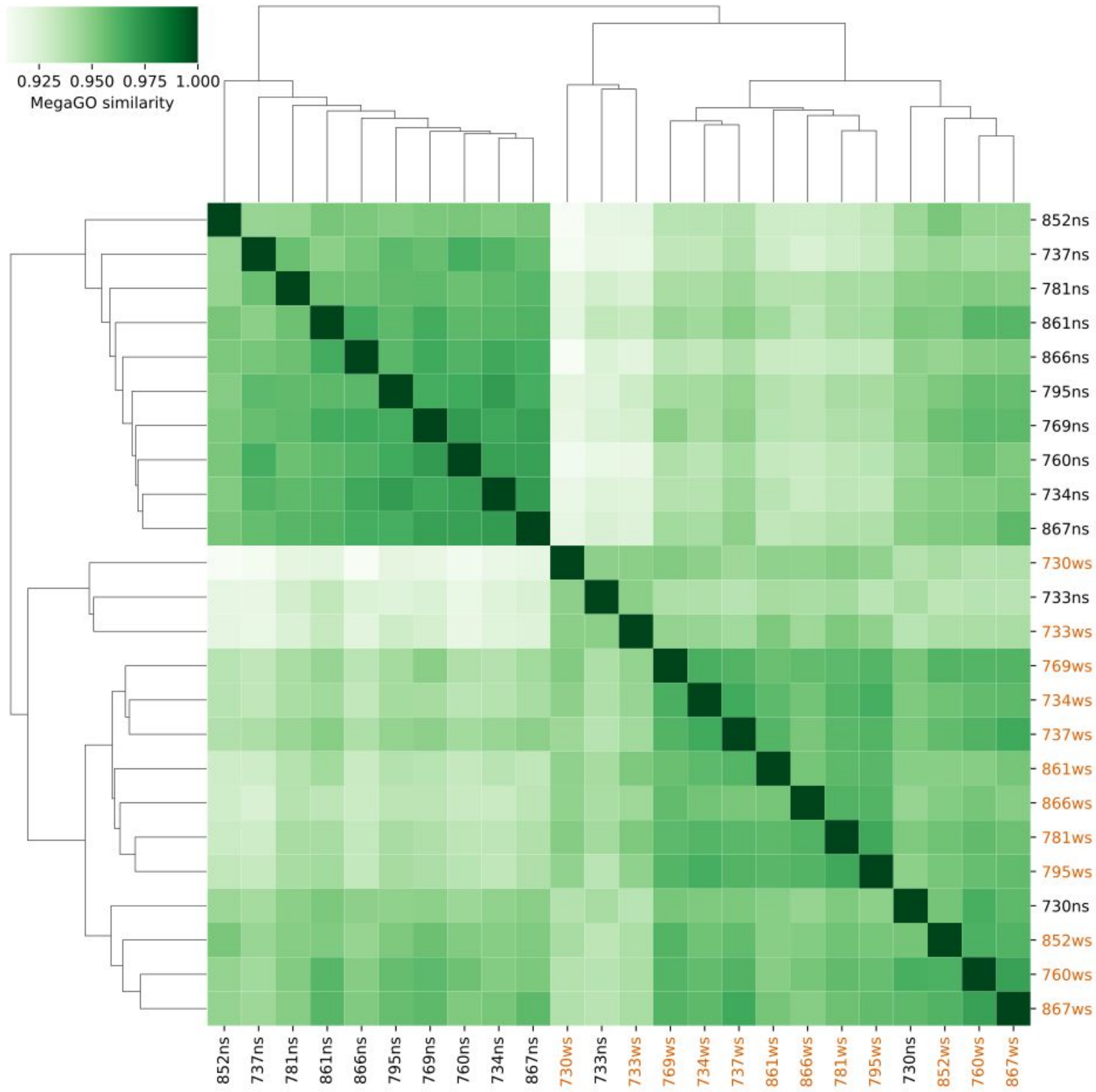


**Supplementary Figure 1**: MegaGO similarities for GO domain molecular function.

### Supplementary Figure 2

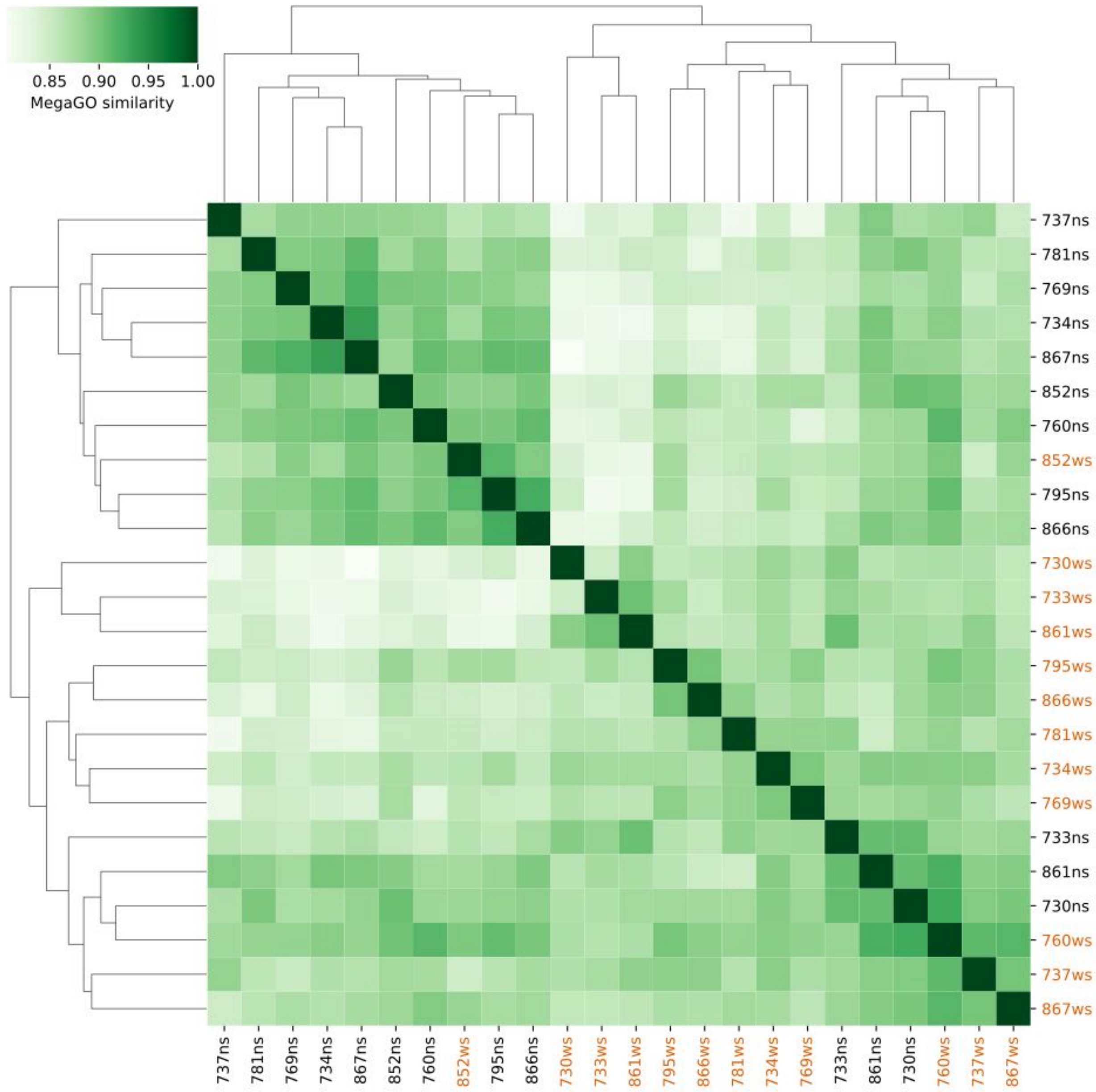


**Supplementary Figure 2**: MegaGO similarities for GO domain cellular component.
